# Generative interpolation and restoration of images using deep learning for improved 3D tissue mapping

**DOI:** 10.1101/2024.03.07.583909

**Authors:** Saurabh Joshi, André Forjaz, Kyu Sang Han, Yu Shen, Vasco Queiroga, Daniel Xenes, Jordan Matelsk, Brock Wester, Arrate Munoz Barrutia, Ashley L. Kiemen, Pei-Hsun Wu, Denis Wirtz

## Abstract

The development of novel imaging platforms has improved our ability to collect and analyze large three-dimensional (3D) biological imaging datasets. Advances in computing have led to an ability to extract complex spatial information from these data, such as the composition, morphology, and interactions of multi-cellular structures, rare events, and integration of multi-modal features combining anatomical, molecular, and transcriptomic (among other) information. Yet, the accuracy of these quantitative results is intrinsically limited by the quality of the input images, which can contain missing or damaged regions, or can be of poor resolution due to mechanical, temporal, or financial constraints. In applications ranging from intact imaging (e.g. light-sheet microscopy and magnetic resonance imaging) to sectioning based platforms (e.g. serial histology and serial section transmission electron microscopy), the quality and resolution of imaging data has become paramount.

Here, we address these challenges by leveraging frame interpolation for large image motion (FILM), a generative AI model originally developed for temporal interpolation, for spatial interpolation of a range of 3D image types. Comparative analysis demonstrates the superiority of FILM over traditional linear interpolation to produce functional synthetic images, due to its ability to better preserve biological information including microanatomical features and cell counts, as well as image quality, such as contrast, variance, and luminance. FILM repairs tissue damages in images and reduces stitching artifacts. We show that FILM can decrease imaging time by synthesizing skipped images. We demonstrate the versatility of our method with a wide range of imaging modalities (histology, tissue-clearing/light-sheet microscopy, magnetic resonance imaging, serial section transmission electron microscopy), species (human, mouse), healthy and diseased tissues (pancreas, lung, brain), staining techniques (IHC, H&E), and pixel resolutions (8 nm, 2 µm, 1mm). Overall, we demonstrate the potential of generative AI in improving the resolution, throughput, and quality of biological image datasets, enabling improved 3D imaging.

## INTRODUCTION

Novel three-dimensional (3D) imaging techniques and algorithms designed to integrate large, multimodal datasets have improved our ability to assess normal anatomy and tissue heterogeneity using anatomical, molecular, -omic probes.^1–7^ Across 3D image modalities, a common challenge emerges: a lack of resolution due to mechanical or financial constraints, or due to the presence of damaged or distorted tissue. Here, we introduce a methodology to repair and enhance 3D biological imaging data using generative artificial intelligence (AI) image interpolation. We demonstrate the utility of this method across serial sectioning-based and intact imaging datasets.

Serial sectioning-based and intact imaging methods both present resolution challenges. Imaging methods that utilize serial sectioning take advantage of the ability to multiplex across tens to hundreds of sections.^2,8,9^ However, sectioning-based techniques face two resolution-limiting hurdles. First, the resolution of the sample is limited by the thickness of the serial sections (4 – 10 µm for histology and ∼40 nm for serial section transmission electron microscopy [ssTEM]). This resolution is further limited during the common practice of intermixing stains (hematoxylin and eosin [H&E], immunohistochemistry [IHC], spatial transcriptomics) at regular intervals.^6,8–12^ Second, the axial resolution of the sample is diminished due to physical artifacts of sectioning, where tissue splitting, folding, and warping can dramatically limit the user’s ability to reconstruct continuous structures.^4,13,14^ In contrast, intact imaging approaches such as magnetic resonance imaging (MRI), computed tomography (CT), and tissue clearing enable 3D views of continuous structures.^15–17^ While the preservation of 3D structure generally enables higher resolution images than serial sectioning approaches, these techniques sacrifice the ability to multiplex across z-planes. Additionally, in spite of the lack of sectioning, resolution problems persist, as the effects of photobleaching, light-sheet absorption, susceptibility to motion artifacts, and signal loss can result in localized loss of tissue connectivity and clarity. ^18–20^

A promising solution lies in the application of generative models and interpolation techniques to enhance the fidelity of reconstructed images. Various generative deep learning models have been employed to synthesize tissue images. Prominent are CycleGANs (Cycle-Consistent Generative Adversarial Networks) and diffusion models.^21–28^ CycleGANs are generative deep learning models that allow for cross modality translation. They have been used for the transformation of H&E-stained slides into synthetic IHC-stained slides that mark specific proteins in tissues.^23–25,29^ Diffusion models have been used to generate magnetic resonance imaging (MRI) and computed tomography (CT) scans to augment the training datasets of deep learning models.^21,27,30^

Despite advances in generative models, limitations persist in achieving synthetic biological images that look realistic, as assessed by rigorous metrics.^21–28,31–33^ Issues such as the accurate representation of subtle or rare textures, cell arrangements, and tissue boundaries are areas of active research.^22,26^ Here, we explore interpolation techniques, such as frame interpolation for large motion (FILM), to enhance the resolution of 3D biological images.^31–34^ Using FILM to generate synthetic intervening slides, we propagate information contained in adjacent slides, which enhances z-axis resolution of 3D microanatomical structures and allows for additional information.

We demonstrate that interpolation of biological images using FILM provides superior performance compared to conventional linear interpolation. FILM-synthesized images can reconstruct microanatomical features, image contrast, and cell counts from damaged slides. Using FILM, 3D reconstructions of semantically segmented synthetic images of complex microanatomical structures - such as ducts and blood vessels - feature fewer artifacts than original, damaged datasets (as assessed considering 13 Haralick features). The versatility of FILM is shown by its applications to different imaging modalities (light microscopy, MRI, ssTEM), species (human, mouse), organs (pancreas, brain, lungs), and pixel resolutions (8 nm, 2 µm, 1mm). These applications highlight the potential of generative AI interpolation techniques such as FILM to enhance spatial resolution, restore and recover damaged image slides, and mitigate information loss in volumetric biomedical imaging.

## MATERIALS AND METHODS

### Specimen acquisition

A sample of non-diseased human pancreas tissue was stained with hematoxylin and eosin (H&E); another similar sample was stained with leukocyte marker CD45 via immunohistochemistry (IHC-CD45). Both samples were from individuals who underwent surgical resection for pancreatic cancer at the Johns Hopkins Hospital.^2^ The H&E-stained dataset consisted of a stack of 101 serially sectioned 4µm apart slides at 5x magnification. H and E are standard histological stains that mark nuclei and cellular structures (H) and ECM (E). The IHC-CD45 stained dataset consisted of 275 slides at 5x magnification where every third slide of the serial section was stained 16µm apart. CD45 is a general marker of leukocytes. This retrospective study was approved by the Johns Hopkins University Institutional Review Board (IRB).

A stack of serial section transmission electron micrographs (ssTEM) within a densely annotated mouse visual cortex petascale image volume (public dataset Minnie65) was obtained through the online Brain Observatory Storage Service and Database (BossDB), created, and managed by the Johns Hopkins Applied Physics Laboratory (APL). This dataset consisted of 100 ssTEM slides captured at a resolution of 8 nm x 8nm x 40 nm.^2,7^

Light-sheet microscopy images of mouse lung were obtained from the Image Data Resource (IDR) public repository.^35,36^ This dataset consisted of 401 serial light-sheet microscopy images captured at a resolution of 3.22µm x 3.22µm x 10µm.

MRI samples of human brain were obtained from the Amsterdam Open MRI Collection (AOMIC).^37^ Specifically, the PIOP2 (Population Imaging of Psychology) cohort consisting of structural MRI scans of students was used. The dataset consisted of 220 structural MRI scans captured at a resolution of 1mm x 1mm x 1mm.

### Segmentation of pancreatic microanatomy in histology slides

CODA, a previously developed semantic segmentation model, was leveraged to segment the H&E-stained pancreas whole slide images (WSIs) into their respective microanatomical components.^2^ CODA was specifically trained for the segmentation of microanatomical components of the pancreas and labeled seven components at a resolution of 2 µm per pixel, including islet of Langerhans, ductal epithelium, blood vessels, fat, acini, extracellular matrix (ECM), and pancreatic intraepithelial neoplasia (PanIN), which are precursor lesions of pancreatic cancer.^2^

### Interpolation between 2D images

Spatial interpolation between 2D slides within a stack was carried out using Frame Interpolation for Large Image Motion (FILM), a model previously developed for temporal interpolation between frames of videos by Reda *et al*.^34^ The model uses a three-step process to generate intermediate frames between two input images: a feature extraction pyramid, optical flow estimation, and feature fusion and frame generation.

The feature extraction pyramid consists of six convolutional layers responsible for extracting features from the input images, each with increasing kernel size and decreasing stride capturing progressively larger receptive fields, extracting features from coarser to finer scales. This coupled with the use of shared weights across scales, allows the model to extract features for both small and large motions efficiently.

The features extracted are then fed into a bi-directional optical flow estimation module. This module calculates the pixel-wise motion vectors (or “flows”) between the features of two input images at each pyramid level. These flows represent the transformation needed to warp the features from one frame to the other. The bi-directional approach allows the model to capture both forward and backward motion, leading to more accurate and detailed interpolations.^34^

With the extracted features and estimated flows, FILM enters the final fusion stage. The aligned features from both input images, along with the flows and the original input images themselves, are concatenated into a single feature pyramid. This captures both the feature information and the motion dynamics between the two frames. Finally, a U-Net decoder architecture processes this fused feature pyramid and generates the final interpolated frame. The U-Net’s skip connections, which bypass several layers within the network and concatenate their outputs directly with the outputs of later layers, ensures that the generated frame retains fine details and maintains consistency with the input images.^34^

FILM used a recursive function (Eq.1) which accounted for the number of input frames, *n*, and the number of recursive passes over which the model would interpolate, *k.* This limited the number of frames that could be generated between the input images to be either one, four, seven, or fifteen frames (Eq.1).

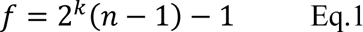

Recognizing the need for flexibility in slide skipping based on user requirements, a time series spanning from 0 to 1 was implemented, with step sizes dynamically determined by the number of skipped slides. This approach generated time points corresponding to the skipped slides, facilitating variable frame interpolation between input pairs.

FILM was pretrained on the Vimeo-90k dataset, a largescale dataset of 89,800 high quality videos designed specifically to train models oriented towards video processing tasks such as frame interpolation, image denoising and resolution enhancement.^34^ The optical flow of this model is already robustly pretrained on a diverse set of videos with different moving objects, such as vehicles, people, and smaller features like cameras and soccer balls. Re-training of the model posed two challenges: a lack of documentation on retraining and perfectly registering histological slides to curate a training dataset. The focus of FILM on optical flow means that the model is sensitive to misalignment in the training images, making histological slides an unfavorable dataset to retrain the optical flow model due to inherent variability in tissue preparation, staining intensities, and sectioning processes, which lead to unpredictable distortions and variations that complicate accurate spatial alignment of a stack of slides.

### Pearson correlation

To characterize the correlation between input image pairs to our model, the Pearson correlation was calculated between pairs of authentic images used to interpolate. This metric allowed for a comparison of three interpolation techniques: nearest-neighbor interpolation, linear interpolation, and FILM. By determining the correlation between the middle interpolated image (furthest from input images) and the corresponding authentic image for each method of interpolation we determined linear interpolation performed the nearest to FILM and hence chose it as the form of interpolation for a more stringent comparison to FILM interpolation (Fig. 2d). The Pearson correlation was calculated using the SciPy stats package available in python.

### Haralick texture features

Thirteen Haralick texture features were calculated to provide a quantitative representation of the texture patterns within an image, offering insights into its spatial arrangements and relationships.^38,39^ The 13 features measured: angular second moment, contrast, correlation, sum of squares variance, inverse difference moment, sum average, sum variance, difference variance, sum entropy, difference entropy, entropy, information measure of correlation 1, and information measure of correlation 2.^38,39^ Contrast measures the intensity variations between neighboring pixels, correlation gauges the linear dependency of gray levels, energy represents the image uniformity, and homogeneity measures the closeness of gray level pairs.

To manage the complexity and high dimensionality of the feature space, dimensionality reduction was carried out using principal component analysis (PCA). PCA transformed the original set of Haralick features into a reduced set of principal components, retaining the most significant information while discarding redundant or less informative aspects. This reduction not only simplifies the interpretation of the data, but also allows for a holistic assessment of image quality, capturing the essential texture information in a more compact form.

Additionally, analysis of the Euclidean distances between authentic and interpolated images was computed using 13 of the Haralick features. By considering the Euclidean distances across all selected Haralick features simultaneously, a comprehensive evaluation of the overall error value was achieved. This validation process ensured that the collective impact of texture features was considered, providing a robust measure of dissimilarity or similarity between images. The combination of Haralick texture features, PCA for dimensionality reduction, and Euclidean distance computation offered a systematic and effective approach for evaluating image quality and texture patterns.

### Cell detection in histological sections

To validate the interpolated IHC images, the CODA cell detection module was used to count the total number of CD45+ cells and compare it with respective authentic images.^2^ For this task, the intensity range of blue pixels was first determined for the nuclei of cells, along with the intensity of brown pixels for positive CD45 stain. Using k-means clustering, the mode blue pixel intensity was determined and selected to represent the hematoxylin channel, while the mode brown pixel intensity was selected to represent the positive stain. With color deconvolution, the cells stained with hematoxylin could be extracted from the remaining tissue, thereby providing a cell count.

### 3D rendering of interpolated 2D images

FILM was used to interpolate stacks of whole slide images (WSIs) of missing or damaged slides, which resulted in the restoration of the whole serial sectioned dataset (Fig. 1b). During post-processing, CODA was used to semantically segment histology slides and MRI images to reconstruct microanatomical tissue structures and whole organs in 3D (Fig. 1b). ^2^ Through manual annotations of microanatomical tissue structures in a small subset of histology slides and whole organ annotations of the brain in a subset of MRI images, CODA allowed for two deep learning models to be trained to recognize these annotations and apply them to the remaining slides/images in the respective datasets, thereby generating stacks of segmented histology slides and MRI images. Labels within the segmented slides/images, corresponding to the annotations could then be used by CODA to reconstruct and visualize 3D tissue structures of interest, such as epithelial ducts in the case of the pancreas, and whole organs such as the brain. Similarly, CODA was leveraged to 3D reconstruct synapses in the mouse brain using pre-segmented ssTEM slides with the appropriate synapse label. Tissue-cleared light-sheet images were separated into their respective RGB channels allowing for three stacks to be obtained, one for each channel. 3D reconstructions of structures within the tissue-cleared light-sheet images of the lung were then generated by creating volumes using stacks of channel-separated images. Specifically, the red channel was used to reconstruct the bronchioles in the mouse lung.

**Fig 1.**
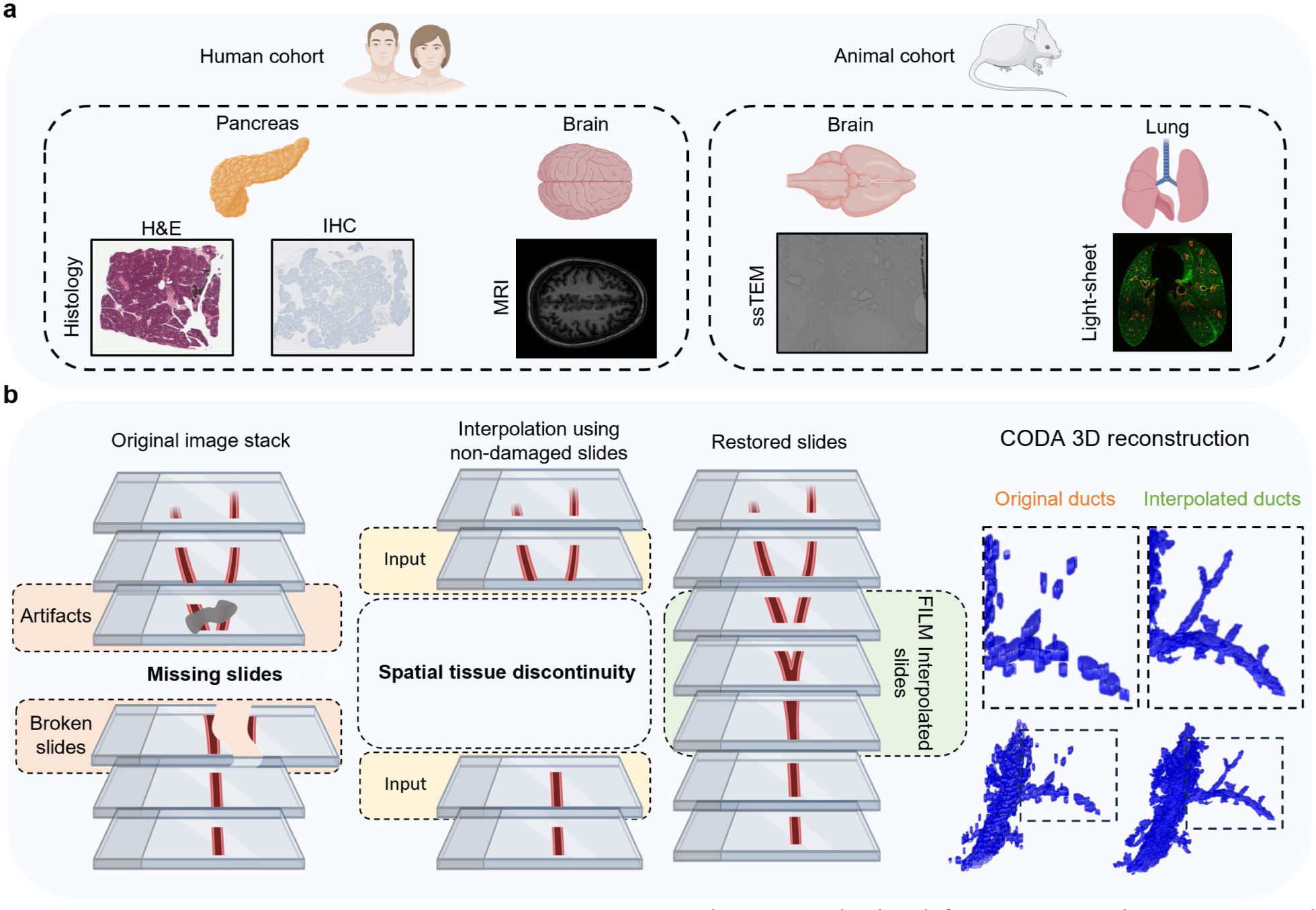
Interpolation workflow and datasets. **(a)** Samples were obtained from two species, mouse, and human. Four different organs were analyzed: human pancreas, human brain, mouse brain, and mouse lung. Five imaging modalities were tested: hematoxylin and eosin (H&E) stained histology slides, immunohistochemistry (IHC) stained histology slides, magnetic resonance imaging (MRI), serial section transmission electron microscopy (ssTEM) slides, and combined tissue clearing with light-sheet microscopy slides. **(b)** Aligned slides are manually searched through to identify missing or damaged slides, and damaged slides are removed from the stack of slides. FILM interpolation is carried out using the sections adjacent to the damaged or missing slides as inputs to recreate slides that were stained differently, missing, or damaged, resulting in a uniform stack of slides. Using CODA, slides are segmented into labeled tissue masks, with each label representing different microanatomical structures in the slide, which is then used to recreate and visualize microanatomical 3D structures in the tissue sample.

**Fig. 2.**
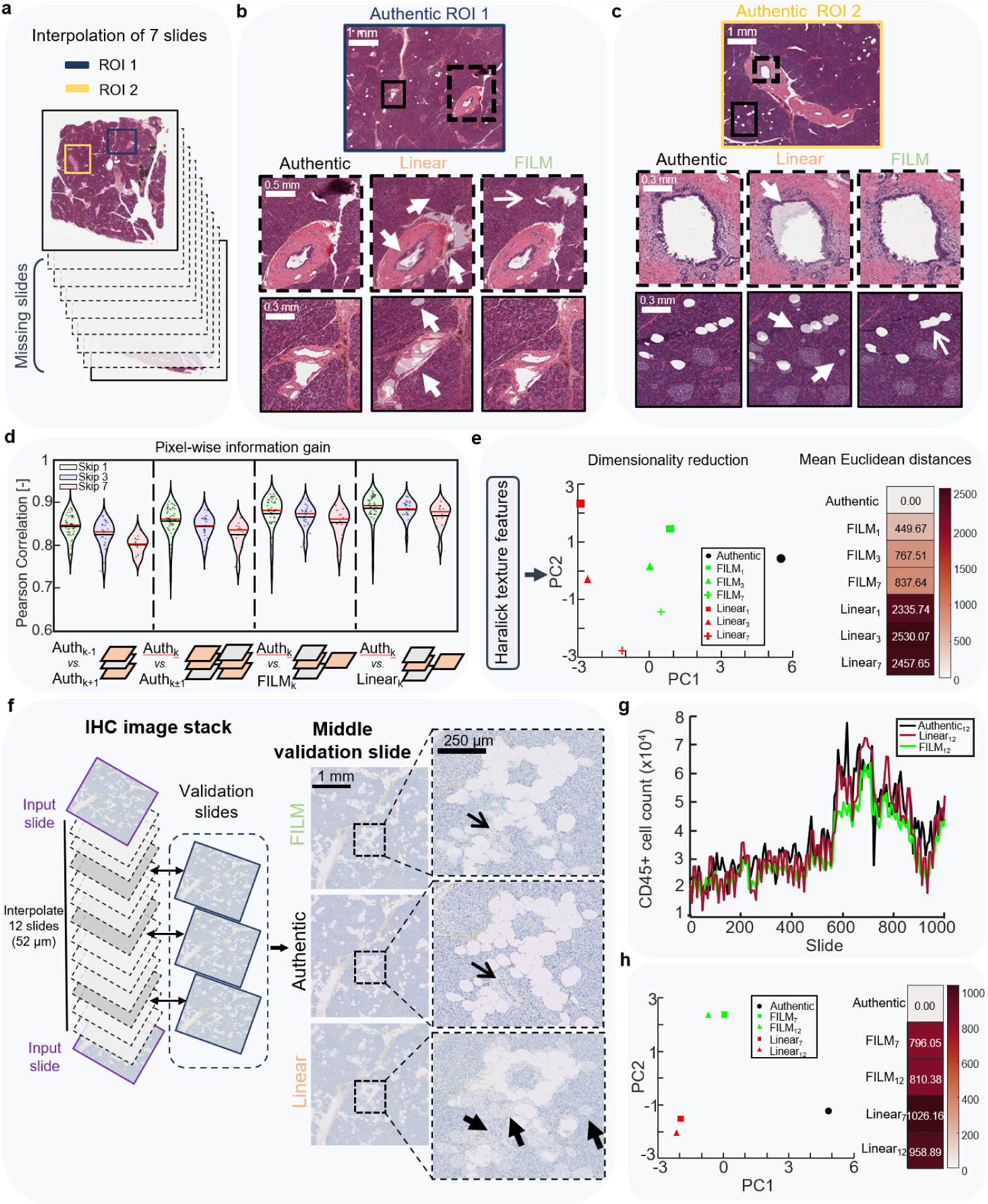
Comparison of linear and FILM interpolations for stacks of histological images of pancreatic tissues. **(a)** Regions of interest (ROI’s) were selected from the whole slide images (WSI’s), ensuring that all microanatomical features (islets of Langerhans, ductal epithelium, blood vessels, fat cells, acini, extra-cellular matrix (ECM), whitespace, and pancreatic intraepithelial neoplasia (PanIN) were present in the ROI and slides were interpolated while skipping 7 slides between adjacent sections, thereby generating 7 slides. **(b)** ROI 1: Comparison of linear and FILM interpolation to the authentic ROI for the middle-interpolated image (image 4) for ductal epithelium and blood vessels. Arrowheads show linear interpolation replacing damage with acini as opposed to whitespace, creating noise around the epithelium layer of the duct, incorrectly generating fat regions, and unable to preserve vessel structure. The arrow shows FILM correctly replaces damage with whitespace. **(c)** ROI 2: Comparison of linear and FILM interpolation to the authentic ROI for the middle-interpolated image (image 4) for ductal epithelium, fat cells, and blood vessels. Arrowheads show linear interpolation creates duct lumen shadows and fat shadows resembling islets as well as non-existent fat regions. **(d)** Pearson correlation compares the correlation between the authentic input images and the nearest-neighbor-interpolated, FILM-interpolated, and linearly interpolated images. **(e)** Principal component analysis of thirteen Haralick features for authentic, FILM, and linearly interpolated images for various numbers of skipped images. Mean Euclidean distance of interpolated images from authentic images based on thirteen Haralick features. **(f)** IHC pancreas slides used to interpolate with authentic slides for validation to compare interpolated images to authentic images. The middle validation slide is visualized for comparison with the interpolated images. Arrow shows how FILM preserves vessel structure, unlike linear interpolation, which was also unable to preserve fat domains (arrowheads). **(g)** Comparison of CD45+ cell counts in authentic images and interpolated images when skipping 12 slides. **(h)** Principal component analysis of thirteen Haralick features for authentic, FILM, and linearly interpolated IHC images for various numbers of skipped images. Mean Euclidean distance of interpolated images from authentic images based on thirteen Haralick features.

### Computing hardware and software

We used Python (v3.8.16) and Tensorflow (2.10.0) for all image interpolations and analysis. For the CODA quantifications and 3D renderings, we used MATLAB (2023a).

For smaller sized images, computers equipped with a single NVIDIA RTX 3090 GPU could easily interpolate them. For larger whole slide images, with dimensions exceeding 14000퀇10000 pixels, using more GPU power would allow to speed up the interpolation processing times. To handle these larger images with higher magnifications, we utilized the Rockfish cluster at Johns Hopkins University, which is equipped with nodes containing four NVIDIA A100 GPUs each. This high-performance computing resource enabled us to interpolate whole slide histological images in shorter times. In case of no access to GPU clusters, users may opt for a tile and stitch approach provided in our code, which allows for tiling of large WSIs, interpolating the tiles individually, and then stitching them back together into WSIs during post-processing.

## RESULTS

### Multi-modal tissue cohorts and interpolation workflows

Here we applied a method based on optical flow, FILM, to restore damages in stacks of 2D images to recover lost microanatomical features in 3D reconstructions of tissue architecture and tissue/cellular composition (Fig. 1).^27^ We procured and tested FILM for a non-diseased pancreatic tissue cohort (stained with H&E and IHC), a structural MRI dataset of the human brain, a stack of ssTEM micrographs of thin sections of the mouse brain, and a mouse lung tissue cleared and imaged under light-sheet microscopy. The selection of these datasets encompassed different image characteristics (size and resolution), species (human, mouse), tissue types (pancreas, brain, lung), imaging modalities (histology, ssTEM, structural MRI, tissue clearing for light-sheet microscopy), and magnifications. This diversity of datasets ensured that the robustness of FILM was evaluated across a broad spectrum of imaging modalities.

FILM, which we compare to other interpolation methods, uses pairs of undamaged 2D images from an image stack to improve spatial resolution or recover lost microanatomical information (Fig. 1b). The user specifies the number of images to be interpolated based on the number of damaged or missing images between the input slides. Using the output interpolated 2D image stacks, 3D volumes can be reconstructed without missing or damaged images (Fig. 1b). This results in improved spatial resolution and reconstruction of tissue components in 3D (Fig. 1b).

### FILM interpolation for stacks of histological slides

We first tested the ability of FILM to interpolate whole slide images from a stack of histological images from human pancreatic tissue samples. Histological slides are often lost or damaged due to improper storage or documentation.^13,14^ The ability of FILM to interpolate slides was compared to a linear interpolation of the same slides and then compared to the corresponding authentic slide (Fig. 2).^32,40–42^ To qualitatively compare the interpolated slides, two ROI’s from the 101 serially sectioned and H&E stained human pancreas dataset were selected based on the tissue structures present. ROIs had a total of eight tissue components, including islets of Langerhans, ductal epithelium, blood vessels, fat, acini, ECM, whitespace, and PanIN (precursor) lesions. Pairs of images were selected one every 8 images (skip 7) of the original stack of authentic images, and the missing 7 images were interpolated (Fig. 2a). Interpolated images were validated against their respective authentic images (Fig. 2, b and c).

We examined ducts and blood vessels due to their complex branching character within the first ROI (Fig. 2b). The authentic image of the duct showed damage fixed by FILM interpolation (top row, top arrow, Fig. 2b). In contrast, the epithelium layer of the duct showed significant noise in the linearly interpolated image due to pixel averaging (top row, bottom arrowhead, Fig. 2b). This caused overlay artifacts absent in FILM, which tracked pixel movements using optical flow for a sharper image. We also observed that linear interpolation replaced the damaged areas with acinus, unlike the whitespace in the authentic slide (top row, top arrowhead, Fig. 2b). In contrast, FILM successfully removed the damage and preserved the whitespace (top row, top arrow, Fig. 2b). Furthermore, FILM preserved the central structure of the duct, whereas linear interpolation thinned and elongated the lumen (top row, middle arrowhead, Fig. 2b). The superiority of FILM over linear interpolation was further seen in the blood vessel microanatomical structures (bottom row, bottom arrowhead, Fig. 2b). With linear interpolation, overlay artifacts were present throughout the entire structure of the blood vessel (bottom arrow, Fig. 2b). Critically, linear interpolation could not preserve the structure of the blood vessel, unlike FILM (Fig. 2b). Linear interpolation also incorrectly generated fat regions absent in the authentic images (bottom row, top arrowhead, Fig. 2b).

In the second ROI, enriched in ducts, fat, and islets, linear interpolation created a duct lumen shadow (top row, top arrowhead, Fig. 2c). In contrast, FILM accurately interpolated the duct without artifact (top row, Fig. 2c). Other key structures were fat and islets, which typically presented a small and faint morphology (bottom row, Fig. 2c). The authentic slide contained 8 fat and 5 islets structures, however linear interpolated images showed fat shadows where the real fat was located (bottom row, top arrowhead, Fig. 2c). Additionally, it generated a non-existent fat region (bottom row, bottom arrowhead Fig. 2c). These fat shadows could be wrongly interpreted as islets, especially in regions where islets are present (bottom row Fig. 2c). Although FILM struggled with overlapping fat, it properly interpolated distinct fat without artifacts and could clearly distinguish islets from fat.

We quantified differences between FILM and linear interpolation of whole slide images using Pearson correlation for each of our scenarios (when skipping 1, 3, and 7 slides) (Fig. 2d). The correlation was calculated (i) between the two input WSIs to the model as well as (ii) between the input WSIs and middle authentic WSIs for each scenario. This correlation (ii) represented the correlation achieved when interpolating images using the nearest neighbor form of interpolation. Lastly, the correlation (iii) between the middle FILM, (iv) the middle linear interpolated image and the middle authentic image for each scenario was calculated. FILM-interpolated WSIs were clearly more correlated to their authentic counterparts than the nearest neighbor-interpolated images. Linearly interpolated images closely matched the correlation obtained between FILM interpolated images and authentic images (Fig. 2d). Hence, linear interpolation was chosen as the benchmark comparative form of interpolation to FILM.

Thirteen Haralick features (angular second moment, contrast, correlation, sum of squares variance, inverse difference moment, sum average, sum variance, difference variance, sum entropy, difference entropy, entropy, information measure of correlation 1, and information measure of correlation 2) were measured to evaluate the interpolated images.^38,39^ The results of each score were averaged for the different tested scenarios (authentic, FILM_skip1_, FILM_skip3_, FILM_skip7_, linear_skip1_, linear_skip3_, and linear_skip7_) (Table S1.), which allowed for principal component analysis (PCA) to be carried out (Fig. 2e). This analysis demonstrated that the FILM-interpolated slides represented more closely the information in the authentic slides, even when skipping seven slides, as compared to linear interpolation. The averaged values were also used to compute the Euclidean distance of the 13 Haralick features between authentic and interpolated images (Fig. 2e). Even skipping 7 slides, FILM images were <1/2 the distance of linear images skipping just 1 slide from authentic images.

Standard metrics, such as mean square error (MSE), structural similarity index measure (SSIM), peak signal-to-noise ratio (PSNR), Spearman correlation, Jaccard correlation, Sobel filter, and channel wise pixel-to-pixel intensity correlation could not quantify the structural errors in microanatomical features from linear interpolation (Fig. 2, b and c). The dominant, easily interpolated acini surrounding microanatomy resulted in similar metric values for linear and FILM, since these metrics are less sensitive to small-pixel deviations compared to large-pixel deviations. Masking out acini to consider only the pixels associated with microanatomical structures was attempted, but registration differences between authentic and interpolated images meant that these metrics only highlighted alignment differences rather than the quality of interpolation.

In sum, FILM can accurately interpolate damaged or missing H&E-stained histological images, which restores lost information in 2D image stacks and, consequently, improves connectivity of microanatomical structures in 3D (see also more below). Unlike linear interpolation, FILM does not generate non-existent microanatomical structures like ducts or fat.

### FILM interpolation for stacks of images stained via immunohistochemistry (IHC)

To further demonstrate the ability of FILM to interpolate histological WSIs, a second human pancreas sample was immunostained (IHC) for leukocyte marker CD45. We note the substantial z-directional distance of 52μm between input slides, equivalent to omitting twelve successive 4-μm-thick sections (Fig. 2f). The target images, for which authentic validation slides were available for comparison, are shaded in dark grey, while the missing slides between the input and target slides are shaded in light grey (Fig. 2f).

We compared the middle target slide interpolated to the middle authentic validation slide using both linear and FILM models (Fig. 2f). When zooming in to a specific fat dense region, linear interpolation artifacts were evident, while FILM lacked such artifacts (bottom row, middle arrowhead, top row, arrow Fig. 2f). Additionally, whereas cells were distinctly observed in the authentic image, the linearly interpolated image showed faintly stained cells covered with white hues resembling fat (bottom row, right arrowhead, Fig. 2f). In contrast, FILM could interpolate distinct cells around the fat and even preserved most of the ductal and ECM structures (top row, arrow, Fig. 4a), unlike the linear model (bottom row, left arrowhead, Fig. 2f).

Using CODA, the total cell count of CD45 positive cells was determined for each of the linear and FILM interpolated images and compared to the cell counts in the authentic slides while skipping and interpolating 12 slides. Linear interpolation resulted in slides with inconsistent CD45+ cell counts which were either much less or greater than those in authentic slides. Conversely, FILM interpolated slides resulted in cell counts which closely matched the cell count trend in authentic slides (Fig 2g). Linearly interpolated slides had a higher percent error in cell count reaching over 90% for certain slides whereas FILM interpolated slides never exceeded 45% error in cell count (Extended Fig. 2f).

Thirteen Haralick texture features were evaluated for authentic and interpolated slides when interpolating 7 and 12 slides. The results of each score were averaged for the different scenarios assessed (authentic, FILM_skip7_, FILM_skip12_, linear_skip7_, and linear_skip12_) (Table S1.). PCA showed FILM-interpolated slides more closely represented authentic slide information along principal component 1, while linear along component 2 (Fig. 2h). The Euclidean distance between authentic and interpolated images demonstrated FILM_skip12_ more closely represented the authentic slides compared to linear_skip7_ (Fig. 2h).

In summary, by interpolating IHC-CD45 stained images and determining the difference in cell count between authentic and interpolated images, we show the ability of FILM to interpolate not only multicellular structures (ducts, blood vessels) in stacks of histological images, but also smaller features such as individual cells.

### FILM interpolation for stacks of MRI and light-sheet microscopy images

Next, we tested the ability of FILM to interpolate images within stacks of MRI images. MRI imaging faces inherent limitations, such as susceptibility to motion artifacts due to prolonged scan times, leading to patient discomfort, and potential for signal loss due to magnetic field in homogeneity that can impact the quality of acquired images. Pairs of images were selected one every 8 images (skip 7) of the original stack of authentic images, and the missing 7 images were interpolated (Fig. 3a). Interpolated images were validated against their respective authentic images (Fig. 3b).

**Fig. 3.**
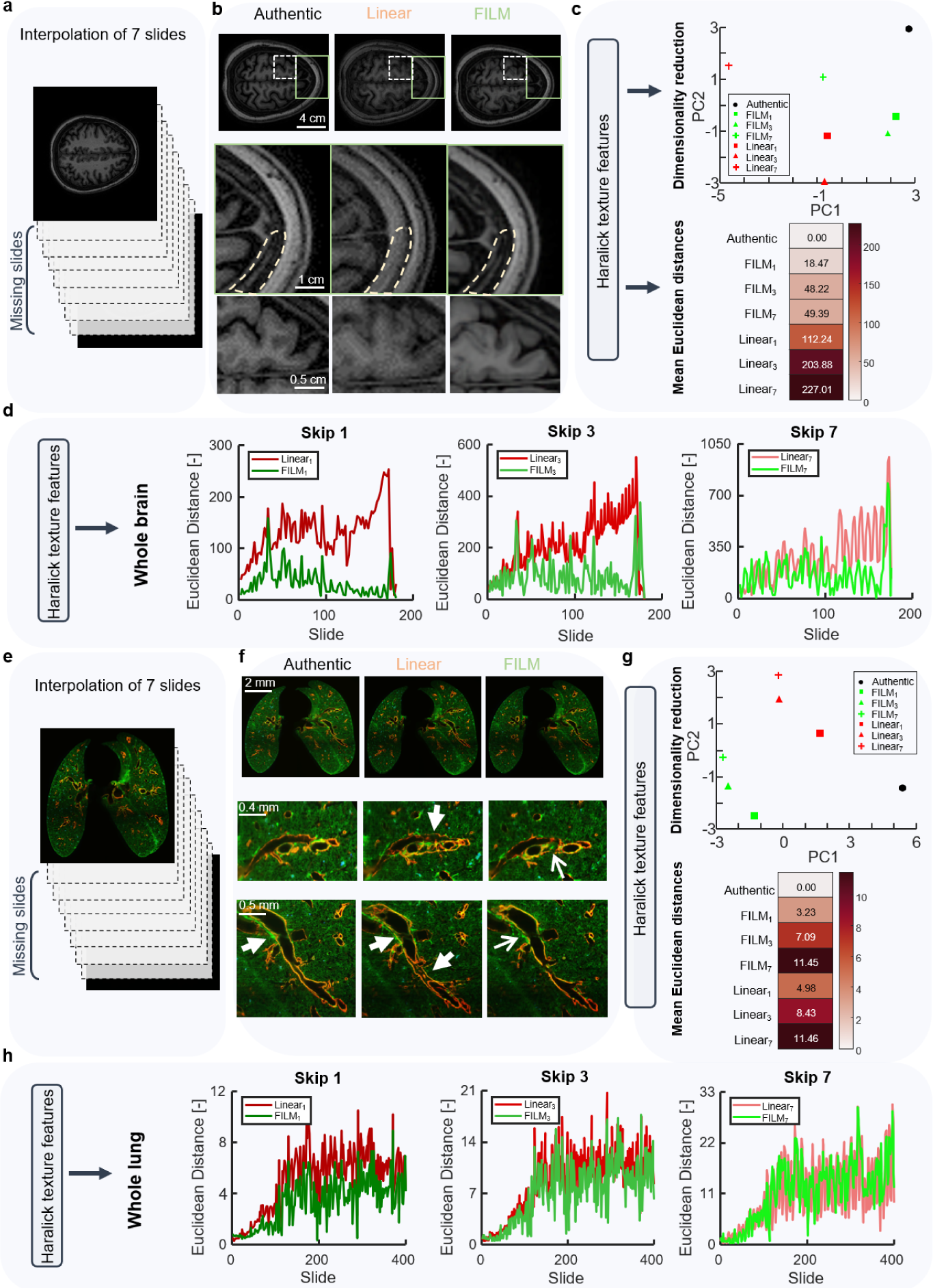
FILM interpolation for stacks of MRI and light-sheet microscopy images. **(a)** MRI images were interpolated while skipping 7 slides between adjacent sections, thereby generating 7 slides. **(b)** Qualitative comparison of linear and FILM interpolation to the authentic image for the middle-interpolated MRI image (image 4). The circled region shows linear interpolation creates band artifacts, unlike FILM. **(c)** Principal component analysis of thirteen Haralick features for authentic, FILM, and linearly interpolated MRI images for various numbers of skipped images. Mean Euclidean distance of interpolated images from authentic images based on thirteen Haralick features. **(d)** Euclidean distance by slide of interpolated images from authentic images based on thirteen Haralick features for various numbers of skipped MRI images. **(e)** Tissue-cleared light-sheet images were interpolated skipping 7 slides between adjacent sections, thereby generating 7 slides. **(f)** Qualitative comparison of linear and FILM interpolations to the authentic image for the middle-interpolated light-sheet image (image 4). Arrowhead shows linear interpolation creates double boundary lines around bronchioles. In second row, the arrowhead shows photobleaching in authentic reduced by linear interpolation and completely removed by FILM (arrow). **(g)** Principal component analysis of thirteen Haralick features for authentic, FILM, and linearly interpolated light-sheet images for various numbers of skipped light-sheet images. Mean Euclidean distance of interpolated images from authentic images based on thirteen Haralick features. **(h)** Euclidean distance by slide of interpolated images from authentic images based on thirteen Haralick features for various numbers of skipped light-sheet images.

Linear interpolation of MRI images caused band artifacts generated around the boundary of the soft tissue, unlike FILM which did not such artifacts (middle row, Fig. 3b). Additionally, FILM could accurately interpolate the soft tissue structure to make biologically accurate structures, whereas linear interpolation created what resembles a grey smudge with significant overlay artifacts (bottom row, Fig. 3b).

To further demonstrate its versatility, we applied FILM to interpolate images within a stack of light-sheet micrographs obtained from a cleared mouse lung. Light-sheet microscopy presents challenges, including photobleaching and light sheet absorption, which may result in uneven illumination, and tissue movement during imaging, which can introduce distortions. Again, pairs of images were selected every 8 images of the authentic stack (Fig. 3e), and interpolated images were compared to their authentic counterpart.

Linear interpolation of light-sheet micrographs created double boundary lines around the bronchioles creating a structure that is biologically inaccurate (middle row, top arrowhead, Fig. 3f) (bottom row, bottom arrowhead, Fig. 3f). In contrast, FILM correctly interpolated the structure of bronchioles to accurately depict the structure observed in the authentic image (middle row, arrow Fig. 3f). In the second row of zoom-ins, we can see that the authentic image suffers from artifacts of light-sheet absorption and photobleaching on the top left side of the bronchiole, which cause bleeding of the green and red channels into the bronchiole (bottom row, arrowhead, Fig. 3f). Linear interpolation reduced these artifacts, but could not remove them entirely (bottom row, top arrowhead Fig. 3f), whereas FILM removed the bleed of the red and green channels (bottom row, arrow Fig. 3f).

For both MRI and light-sheet microscopy datasets, thirteen Haralick texture features introduced above were measured to compare authentic and interpolated images, when interpolating 1, 3, and 7 slides. The results for each score were averaged for the different comparisons (authentic, FILM_skip1_, FILM_skip3_, FILM_skip7_, linear_skip1_, linear_skip3_, and linear_skip7_) (Table S1.), and shown in a principal component analysis (PCA) plane (Fig 3, c and g). For MRI images, FILM-interpolated slides represented more closely the information in the authentic slides, even when skipping seven slides, compared to linear interpolation. The averaged values were also used to compute the Euclidean distance between authentic and interpolated images (Fig 3c). Again, even when skipping seven slides, FILM-interpolated slides were < 1/2 the Euclidean distance between the authentic slides and the linearly interpolated slides when skipping only one slide. The Euclidean distance by slide further emphasizes the superiority of FILM over linear interpolation as the Euclidean distance increased when progressing through the stack of slides and interpolating linearly as opposed to FILM (Fig 3d).

Similarly, the PCA analysis for light-sheet interpolated images showed that FILM interpolated images were more closely representative of the authentic images for principal component 2, while the linear interpolated images were closer for principal component 1 (Fig 3g). Nevertheless, when considering the mean Euclidean distance, FILM outperformed linear interpolation for each individual skip scenario (Fig 3g). The Euclidean distance by slide increased when interpolating linearly through the stack as opposed to FILM, for which it remained consistent through the stack. (Fig 3h).

In sum, we demonstrated the ability of FILM to interpolate MRI and light-sheet images more accurately than linear interpolation. FILM reduces motion artifacts in MRI images whereas linear interpolation exaggerates these artifacts, resulting in band artifacts. FILM reduces photobleaching and light-sheet absorption artifacts present in the authentic light-sheet images, whereas linear interpolation cannot. Elimination of such artifacts allows for more accurate 3D reconstructions of whole organ structures and microanatomical structures in tissue samples.

### FILM interpolation and restoration of ssTEM images

A stack of serial section transmission electron micrographs (ssTEM) of the mouse brain was used to show our ability to interpolate not only histological sections, but also EM micrographs of tissue sections (Fig. 4a). Authentic tiles shown represent a 2000퀇3500 pixel tile of the authentic whole slide image (Fig. 4a). Thick irregular black lines were observed across most of the slides in the authentic stack of images, which correspond to damage due to unavoidable tissue tear during processing of thin sections (left column, arrows, Fig. 4b). For a randomly selected subset of 100 continuous slides from a stack of the >13,000 ssTEM slides, we found that >70% were damaged, many of them containing >1 damaged region. Additionally, fainter grey lines were observed, going horizontally across the authentic images, which are artifacts of image stitching (top row, bound by red box Fig. 4b). Interpolation between two undamaged EM slides using FILM could not only remove the damage to the slides while preserving the structures within them, but also significantly reduce stitching artifacts (right column, Fig. 4b).

**Fig. 4.**
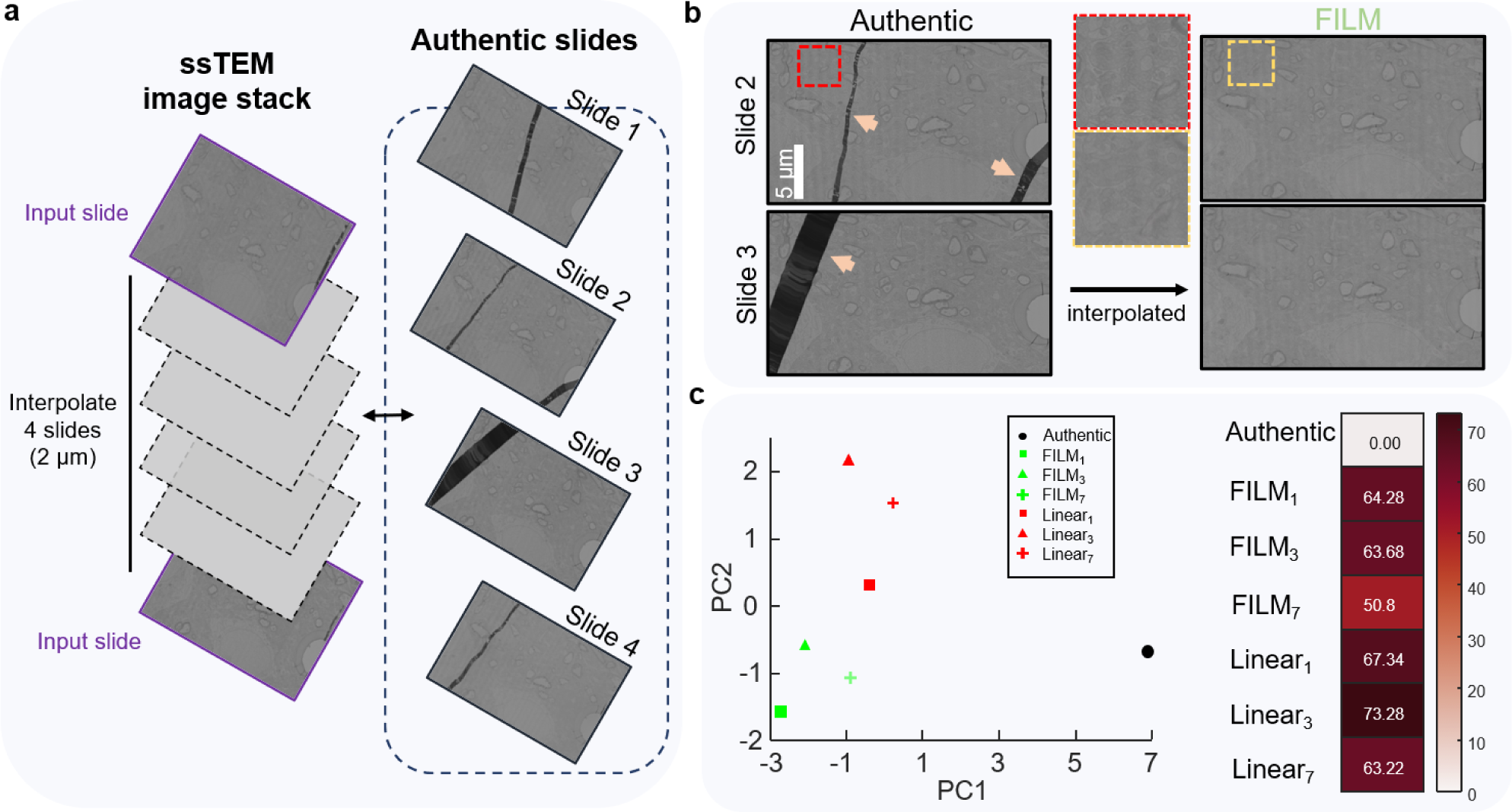
FILM interpolation for a stack of ssTEM images. **(a)** ssTEM slides were interpolated while skipping 4 slides between adjacent sections, thereby generating 4 slides. **(b)** FILM interpolation of mouse brain ssTEM slides to remove damage from slides (arrowheads) and reduce stitching artifacts (red box). **(c)** Principal component analysis of thirteen Haralick features for authentic, FILM, and linearly interpolated ssTEM images for various skipped images. Mean Euclidean distance of interpolated images from authentic images based on thirteen Haralick features.

Thirteen Haralick features were measured for the authentic and interpolated images when interpolating 1, 3, and 7 slides. The results of each score were averaged for the different tested scenarios (authentic, FILM_skip1_, FILM_skip3_, FILM_skip7_, linear_skip1_, linear_skip3_, and linear_skip7_) (Table S1.). PCA showed that FILM-interpolated slides more closely represented authentic slide information along principal component 2, while linear along component 1 (Fig. 4c). Euclidean distance between authentic and interpolated images where it can be seen that FILM_skip1_ more closely represents the authentic than linear_skip1_ and similarly for the instance of skipping 3 and 7 slides (Fig. 4c).

In sum, we demonstrate the ability of FILM to eliminate damage in ssTEM slides. This allows for more accurate 3D reconstructions of the neural pathways by decreasing loss connectivity which arises due to the damage on individual 2D sections.

### 3D reconstruction of FILM-interpolated images

To better show the application of FILM to enhance 3D visualization of microanatomical structures from 2D interpolated images, FILM interpolation was applied across different image modalities such as histological (H&E and IHC), MRI, ssTEM, and light-sheet images. Sematic segmentation and subsequent concatenation of the 2D segmented images into a volume allowed visualization of microanatomical features in three dimensions.

Using CODA, we 3D reconstructed the epithelial duct from the pancreatic H&E dataset (Fig. 5a). The 3D reconstruction of the authentic volume skipping 7 images demonstrates the loss in ductal connectivity as a result of missing or damaged slides. Linear interpolation of the H&E samples created noise around the structure of the duct when 3D reconstructed and was unable to preserve the branching structure of the duct (zoom-in, Fig. 5a). On the other hand, FILM was able to restore the microanatomical connectivity in the 3D reconstruction of the main and smaller branches of the duct, while also creating a smoother volume without the propagation of noise (Supplementary Video 1).

**Fig. 5.**
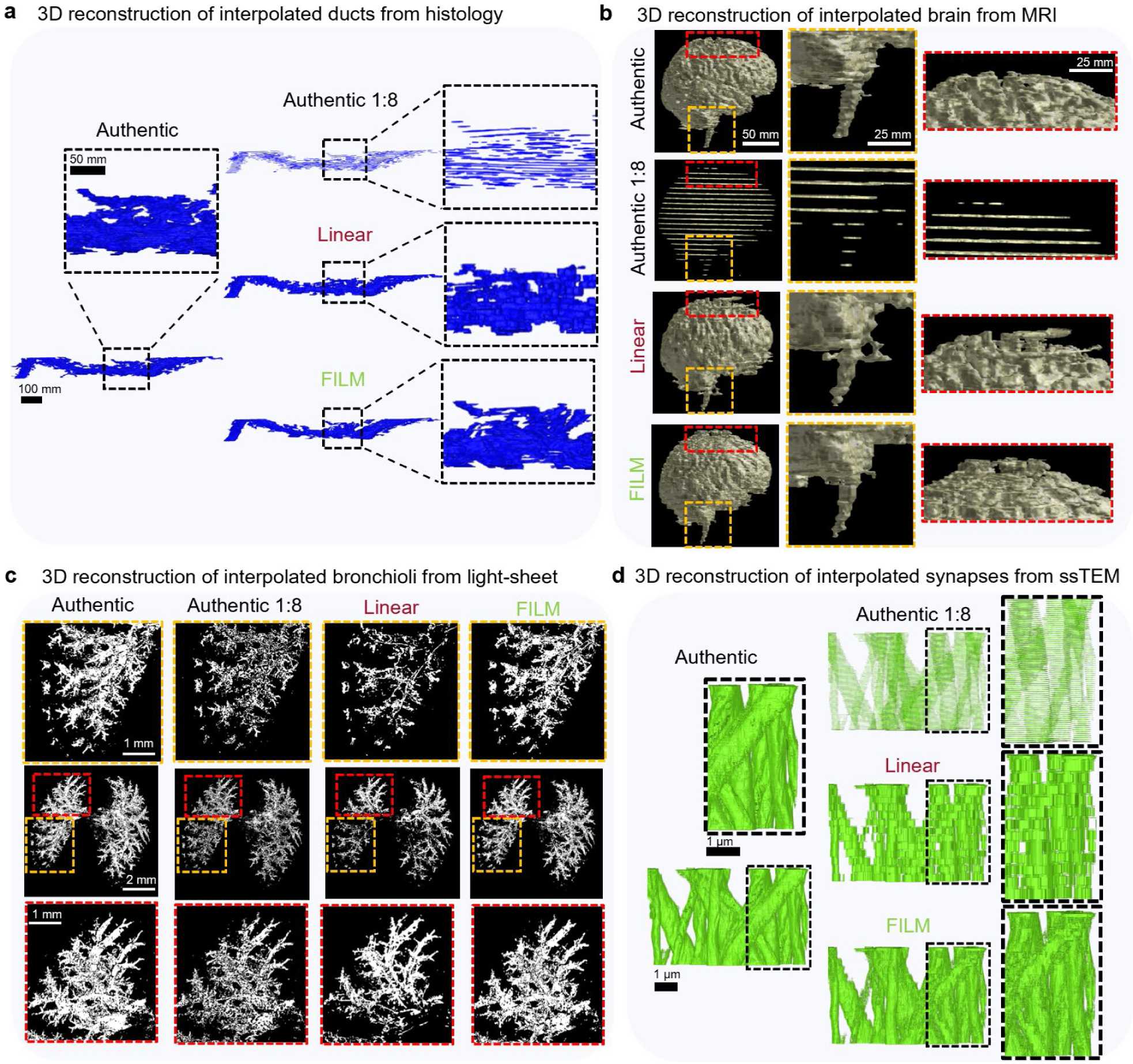
3D reconstruction of interpolated images. **(a)** Comparison of 3D reconstructions of pancreatic duct when skipping 7 slides between authentic images and when interpolating the missing slides using linear and FILM interpolations. **(b)** Comparison of 3D reconstructions of brain MRI images when skipping 7 images between authentic images and when interpolating the missing slides using linear and FILM interpolations. **(c)** Comparison of 3D reconstructions of bronchioles from light-sheet images of the mouse lung when skipping 7 images between authentic images and when interpolating the missing slides using linear and FILM interpolations. **(d)** Comparison of 3D reconstructions of synapses from ssTEM slides of the mouse brain when skipping 7 images between authentic images and when interpolating the missing slides using linear and FILM interpolations.

Similarly, CODA was used to 3D reconstruct a whole human brain using the stack of MRI images. A comparison of the authentic volume to the authentic volume skipping 7 images showed how connectivity was lost as a result of damaged or missing images. The authentic volume skipping 7 images also lacked the topographical structure of the brain seen in the authentic volume, replacing the topography with single planes of information (Fig 5b). Using linear interpolation to recover the missing or damaged scans resulted in increased edges in 3D volumes, which resembled objects extruding abnormally out of the brain. This is especially evident around the base of the brain where the brain stem starts and at the top of the brain towards the skull cap (Fig. 5b). When interpolating images using FILM, the 3D reconstructed volume resembled more closely that of the authentic one, with accurate indentations around the surface of the brain and even accurate reconstruction of the branching brain stem structure.

Tissue-cleared light-sheet images were separated by channel and used to 3D reconstruct the bronchioles in a mouse lung (Fig 5c). Similarly, a comparison of the authentic volume to a downsampled reconstruction of the authentic volume (skipping 7 images between adjacent z-planes) demonstrates the loss in connectivity of the bronchioles in 3D as a result of damaged or missing image scans. The use of linear interpolation to recover missing z-planes did not improve the connectivity of the bronchioles in 3D, but rather more closely resembled the structure of the downsampled volume (Fig. 5c). FILM recovered the missing planes, which restored the connectivity of the bronchioles, and consequently resulted in a volume that resembled the authentic biospecimen.

Segmented ssTEM images were interpolated using FILM and linear interpolation to 3D reconstruct synapses in the mouse brain. A qualitative assessment between the authentic volume and downsampled recreation of the authentic volume (skipping 7 images between adjacent z-planes) shows the loss in synapse connectivity (Fig. 5d). Linear interpolation to recover the missing z-planes results in the creation of a low resolution volume with blocked structures. Conversely, using FILM to recover the missing planes resulted in a higher resolution 3D volume, which resembled that of the authentic volume, and allowed for the synapse connectivity to be restored (Supplementary Video 2).

In sum, missing or damaged slides and images in biomedical image stacks cause significant loss in 3D spatial information which hinders the accurate 3D reconstruction of microanatomical and whole organ structures from these 2D image stacks. We demonstrate that linear interpolation is not sufficiently robust to recover the information lost in complex biomedical images, resulting in inaccurate 3D reconstructions. In contrast, the optical flow-based model FILM can recover more information to allow for 3D reconstructions that resemble their authentic counterparts.

## DISCUSSION

We are entering an era in which 3D imaging of biomedical samples has become a requirement as 2D assessments are not sufficient in capturing the content and morphology of multi-cellular structures, rare events, and spatial relationships among different cell types.^1^ Various models have been developed to leverage 2D biomedical image stacks of histology slides, MRI images, ssTEM slides, and tissue-cleared light-sheet images to reconstruct volumes of microanatomical structures and whole organs. Such models rely on the quality of individual 2D images within the image stacks for the accurate reconstruction of volumes. Limitations in z-resolution often arise due to missing slides and images, tissue damage, and the high cost associated with imaging.

Here, we address these challenges by leveraging FILM and its ability to extract and track features in biomedical images using optical flow for image interpolation. By interpolating between undamaged slides to generate missing or damaged slides, we bridge gaps in z-resolution. This technique enhances 3D reconstructions and mitigates issues arising from damaged or missing slides. This method broadens the applicability of 2D biomedical image stacks for 3D reconstructions and quantitative assessments of cellular composition, tissue topography, and degree of branching of ducts and blood vessels in volumetric tissues.

We conducted a thorough comparative assessment of FILM to linear interpolation using thirteen Haralick texture features. Linear interpolation, which averages pixel intensities creating hued colors and structures, cannot create realistic biomedical images. As the number of images skipped increases, linearly interpolated images further degrade in authenticity, especially for the images furthest from input images (middle-interpolated image). For large number of skipped images (skip 7), the middle-interpolated image presents strong hues as pixel intensities deviate largely between input images. Conversely, FILM can interpolate biomedical images that resemble their authentic counterparts.

By interpolating images using FILM, we reduce the time required for image acquisition. This is especially applicable when considering MRI scans and the time spent by patients in the machine, which can lead to patient discomfort and, in extension, motion artifacts that hinder imaging quality. Similarly, for light-sheet microscopy, we demonstrate the ability of FILM to accurately interpolate images in the z-direction reducing the number z-steps required during image acquisition. This significantly decreases the total time required to image a sample as samples are imaged tile by tile laterally before moving onto the next z-level. Collection time increases exponentially with the lateral size of the sample, from minutes for a 10^4^ µm^3^ sample at a spatial resolution of 500 nm to a week for a 10^8^ µm^3^ sample at the same resolution.^16^ FILM interpolation helps address this limitation.

In conclusion, our work goes beyond existing methods of image translation which use CycleGAN’s and diffusion models to generate biomedical images by leveraging FILM’s method of image interpolation. Where image translation would require physical access to the slides of interest to be translated, our workflow interpolates missing or inaccessible slides, restores damaged images, eliminates artifacts of image stitching, and works with a wide range of complex multimodal biomedical images.

## Data availability

The data analyzed here is available from the corresponding author upon request.

## Author contributions

D.W., S.J., A.F, A.L.K., and P.W. conceived the project. D.W., A.L.K., P.W., and A.M.B. supervised the study. A.L.K., and A.F. collected and processed the human pancreas samples. D.X., J.M. and B.W. collected the mouse brain samples. S. J., A.F, collected and processed the mouse lung and human brain samples. S.J., A.F., K.S.H., Y.S., and P.W. conducted the image analysis, quantifications, and validation. S.J., A.F., A.L.K., and D.W. wrote the first draft of the manuscript, which all authors edited and approved.

## Figures and Captions

**Extended Fig 2.**
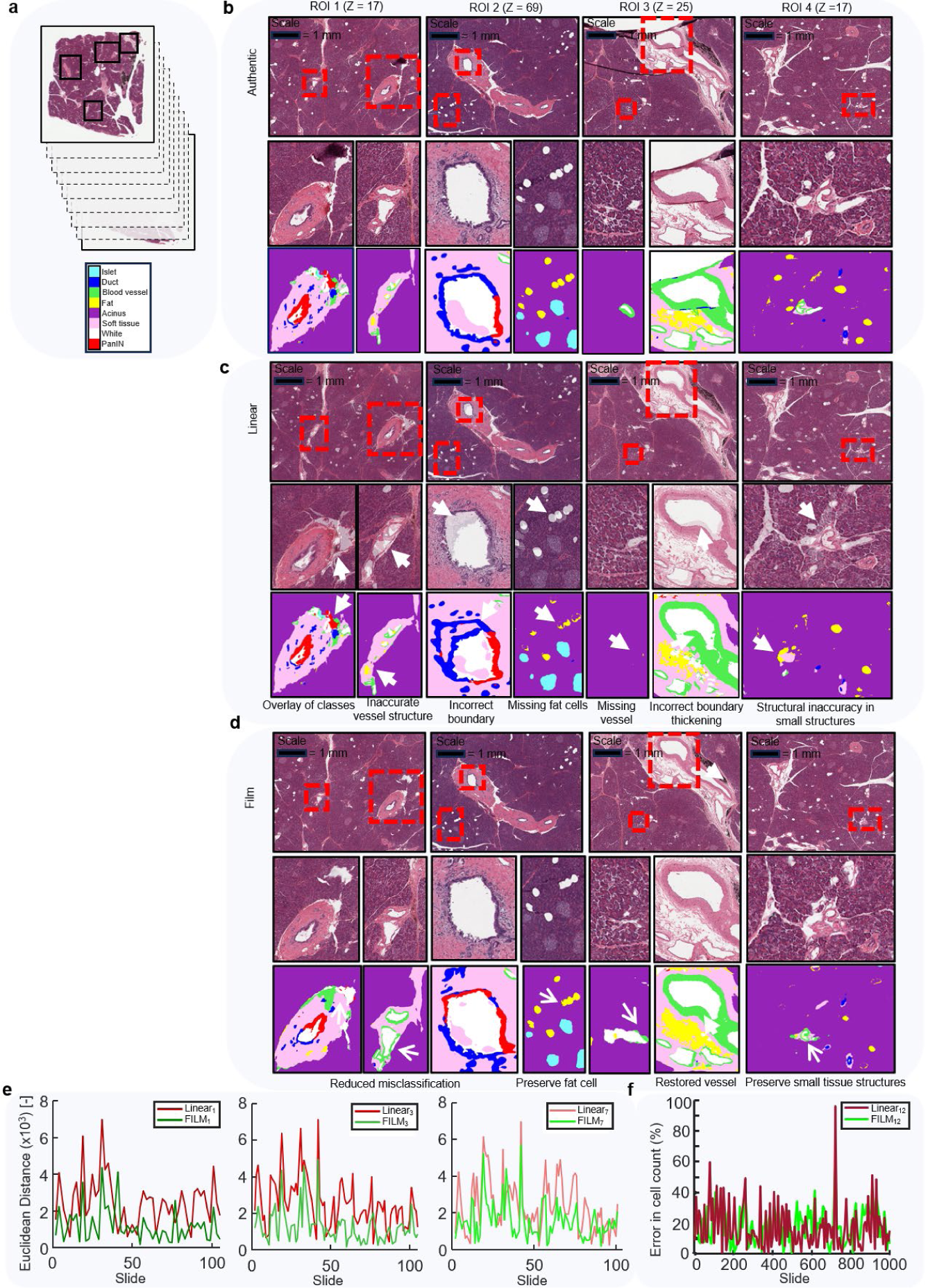
Qualitative comparison of linear and FILM interpolations to authentic H&E-stained histological slides of a human pancreas when skipping 7 slides for four different ROI’s. **(a)** Four ROIs were selected from H&E-stained whole slide images (WSI’s). Slides were interpolated when skipping 7 slides between adjacent sections, thereby generating 7 slides. **(b)** The top row of authentic images shows the middle skipped z-slide of all four different ROIs selected for interpolation. The middle row of zoom-ins of authentic images shows microanatomical structures observed within the different ROI’s. The third row of zoom-ins shows the CODA classification of these microanatomical structures. **(c)** The top row of linearly interpolated images shows the middle interpolated z-slide of all four different ROI’s corresponding to the authentic images. The middle row of zoom-ins of linearly interpolated images shows microanatomical structures generated by linear interpolation within the different ROI’s. The third row of zoom-ins shows the CODA classification of these linearly interpolated microanatomical structures. **(d)** The top row of FILM interpolated images shows the middle interpolated z-slide of all four different ROI’s corresponding to the authentic images. The middle row of zoom-ins of FILM interpolated images shows microanatomical structures generated by FILM within the different ROI’s. The third row of zoom-ins shows the CODA classification of these FILM interpolated microanatomical structures. **(e)** Euclidean distance by slide of interpolated images from authentic images based on thirteen Haralick features for ROI 1 and ROI 2. **(f)** Percent error in CD45+ cell count by slide between authentic and interpolated images when skipping 12 slides.

**Table S1.** Mean Haralick texture feature scores for each dataset and skip scenario.

| HE WSI |  |  |  |  |  |  |  |  |  |  |  |  |  |
| --- | --- | --- | --- | --- | --- | --- | --- | --- | --- | --- | --- | --- | --- |
| Dataset | Energy | Contrast | Correlation | Variance | Homogeneti | Sum Average | Sum Variance | Sum Entropy | Entropy | Difference Variance | Difference Entropy | Info Meas of Corr | Infor Meas of Corr 2 |
| Authentic | 1.42E-02 | 5.62E+02 | 9.57E-01 | 6.51E+03 | 3.32E-01 | 3.08E+02 | 2.55E+04 | 7.34E+00 | 1.12E+01 | 3.39E-04 | 4.82E+00 | -2.65E-01 | 9.83E-01 |
| FILM_1 | 1.23E-02 | 4.11E+02 | 9.68E-01 | 6.43E+03 | 3.30E-01 | 3.07E+02 | 2.53E+04 | 7.31E+00 | 1.11E+01 | 3.48E-04 | 4.68E+00 | -2.76E-01 | 9.85E-01 |
| FILM_3 | 1.19E-02 | 3.92E+02 | 9.69E-01 | 6.36E+03 | 3.28E-01 | 3.08E+02 | 2.51E+04 | 7.32E+00 | 1.11E+01 | 3.45E-04 | 4.67E+00 | -2.76E-01 | 9.85E-01 |
| FILM_7 | 1.17E-02 | 3.91E+02 | 9.69E-01 | 6.35E+03 | 3.26E-01 | 3.08E+02 | 2.50E+04 | 7.35E+00 | 1.11E+01 | 3.40E-04 | 4.68E+00 | -2.77E-01 | 9.85E-01 |
| Linear_1 | 1.34E-02 | 3.15E+02 | 9.74E-01 | 6.05E+03 | 3.31E-01 | 3.08E+02 | 2.39E+04 | 7.29E+00 | 1.10E+01 | 3.51E-04 | 4.59E+00 | -2.84E-01 | 9.86E-01 |
| Linear_3 | 1.29E-02 | 3.32E+02 | 9.72E-01 | 6.01E+03 | 3.28E-01 | 3.08E+02 | 2.37E+04 | 7.31E+00 | 1.10E+01 | 3.44E-04 | 4.62E+00 | -2.82E-01 | 9.86E-01 |
| Linear_7 | 1.26E-02 | 3.63E+02 | 9.70E-01 | 6.02E+03 | 3.25E-01 | 3.08E+02 | 2.37E+04 | 7.36E+00 | 1.11E+01 | 3.36E-04 | 4.67E+00 | -2.82E-01 | 9.86E-01 |
| IHC WSI |  |  |  |  |  |  |  |  |  |  |  |  |  |
| Dataset | Energy | Contrast | Correlation | Variance | Homogeneti | Sum Average | Sum Variance | Sum Entropy | Entropy | Difference Variance | Difference Entropy | Info Meas of Corr | Infor Meas of Corr 2 |
| Authentic | 3.30E-02 | 5.30E+02 | 5.76E-01 | 6.31E+02 | 3.80E-01 | 4.34E+02 | 1.99E+03 | 6.24E+00 | 9.96E+00 | 4.36E-04 | 4.65E+00 | -1.71E-01 | 9.18E-01 |
| FILM_7 | 2.14E-02 | 3.03E+02 | 6.76E-01 | 4.72E+02 | 3.56E-01 | 4.35E+02 | 1.58E+03 | 6.34E+00 | 1.00E+01 | 3.93E-04 | 4.50E+00 | -1.79E-01 | 9.25E-01 |
| FILM_12 | 1.98E-02 | 2.93E+02 | 6.85E-01 | 4.69E+02 | 3.51E-01 | 4.35E+02 | 1.58E+03 | 6.37E+00 | 1.01E+01 | 3.83E-04 | 4.50E+00 | -1.79E-01 | 9.26E-01 |
| Linear_7 | 2.08E-02 | 2.95E+02 | 6.50E-01 | 4.26E+02 | 3.28E-01 | 4.33E+02 | 1.41E+03 | 6.33E+00 | 1.01E+01 | 3.42E-04 | 4.61E+00 | -1.57E-01 | 9.05E-01 |
| Linear_12 | 2.06E-02 | 3.11E+02 | 6.43E-01 | 4.39E+02 | 3.25E-01 | 4.33E+02 | 1.45E+03 | 6.35E+00 | 1.02E+01 | 3.35E-04 | 4.64E+00 | -1.55E-01 | 9.04E-01 |
| MRI |  |  |  |  |  |  |  |  |  |  |  |  |  |
| Dataset | Energy | Contrast | Correlation | Variance | Homogeneti | Sum Average | Sum Variance | Sum Entropy | Entropy | Difference Variance | Difference Entropy | Info Meas of Corr | Infor Meas of Corr 2 |
| Authentic | 4.40E-01 | 7.62E+01 | 9.26E-01 | 5.32E+02 | 7.18E-01 | 2.40E+01 | 2.05E+03 | 3.39E+00 | 4.77E+00 | 3.73E-03 | 2.35E+00 | -4.12E-01 | 9.41E-01 |
| FILM_1 | 4.42E-01 | 6.99E+01 | 9.33E-01 | 5.29E+02 | 7.28E-01 | 2.40E+01 | 2.04E+03 | 3.39E+00 | 4.73E+00 | 3.54E-03 | 2.28E+00 | -4.41E-01 | 9.51E-01 |
| FILM_3 | 4.42E-01 | 6.68E+01 | 9.35E-01 | 5.23E+02 | 7.29E-01 | 2.40E+01 | 2.02E+03 | 3.39E+00 | 4.72E+00 | 3.54E-03 | 2.26E+00 | -4.44E-01 | 9.51E-01 |
| FILM_7 | 4.32E-01 | 6.45E+01 | 9.37E-01 | 5.23E+02 | 7.24E-01 | 2.45E+01 | 2.03E+03 | 3.45E+00 | 4.81E+00 | 3.18E-03 | 2.29E+00 | -4.40E-01 | 9.58E-01 |
| Linear_1 | 4.34E-01 | 5.51E+01 | 9.46E-01 | 5.10E+02 | 7.24E-01 | 2.41E+01 | 1.98E+03 | 3.42E+00 | 4.76E+00 | 3.53E-03 | 2.26E+00 | -4.41E-01 | 9.53E-01 |
| Linear_3 | 4.38E-01 | 5.02E+01 | 9.48E-01 | 4.92E+02 | 7.23E-01 | 2.41E+01 | 1.92E+03 | 3.39E+00 | 4.68E+00 | 4.04E-03 | 2.24E+00 | -4.40E-01 | 9.50E-01 |
| Linear_7 | 4.21E-01 | 5.15E+01 | 9.45E-01 | 4.87E+02 | 7.12E-01 | 2.46E+01 | 1.90E+03 | 3.49E+00 | 4.86E+00 | 3.67E-03 | 2.32E+00 | -4.30E-01 | 9.57E-01 |
| Light-sheet |  |  |  |  |  |  |  |  |  |  |  |  |  |
| Dataset | Energy | Contrast | Correlation | Variance | Homogeneti | Sum Average | Sum Variance | Sum Entropy | Entropy | Difference Variance | Difference Entropy | Info Meas of Corr | Infor Meas of Corr 2 |
| Authentic | 3.89E-01 | 4.25E-01 | 9.87E-01 | 1.94E+01 | 9.22E-01 | 6.25E+00 | 7.70E+01 | 2.61E+00 | 2.85E+00 | 5.36E-03 | 7.05E-01 | -7.19E-01 | 9.67E-01 |
| FILM_1 | 3.97E-01 | 3.80E-01 | 9.88E-01 | 1.88E+01 | 9.29E-01 | 6.25E+00 | 7.46E+01 | 2.57E+00 | 2.79E+00 | 5.57E-03 | 6.58E-01 | -7.49E-01 | 9.72E-01 |
| FILM_3 | 3.98E-01 | 3.57E-01 | 9.89E-01 | 1.80E+01 | 9.29E-01 | 6.24E+00 | 7.16E+01 | 2.56E+00 | 2.78E+00 | 5.66E-03 | 6.52E-01 | -7.48E-01 | 9.72E-01 |
| FILM_7 | 3.96E-01 | 3.39E-01 | 9.89E-01 | 1.71E+01 | 9.29E-01 | 6.25E+00 | 6.81E+01 | 2.57E+00 | 2.78E+00 | 5.88E-03 | 6.53E-01 | -7.45E-01 | 9.72E-01 |
| Linear_1 | 3.89E-01 | 3.42E-01 | 9.88E-01 | 1.84E+01 | 9.26E-01 | 6.24E+00 | 7.32E+01 | 2.60E+00 | 2.82E+00 | 5.95E-03 | 6.77E-01 | -7.26E-01 | 9.67E-01 |
| Linear_3 | 3.88E-01 | 3.15E-01 | 9.89E-01 | 1.77E+01 | 9.28E-01 | 6.25E+00 | 7.05E+01 | 2.60E+00 | 2.81E+00 | 6.72E-03 | 6.63E-01 | -7.31E-01 | 9.69E-01 |
| Linear_7 | 3.86E-01 | 3.10E-01 | 9.88E-01 | 1.71E+01 | 9.27E-01 | 6.25E+00 | 6.81E+01 | 2.60E+00 | 2.81E+00 | 7.22E-03 | 6.63E-01 | -7.31E-01 | 9.69E-01 |
| Cryo-electron microscopy |  |  |  |  |  |  |  |  |  |  |  |  |  |
| Dataset | Energy | Contrast | Correlation | Variance | Homogeneti | Sum Average | Sum Variance | Sum Entropy | Entropy | Difference Variance | Difference Entropy | Info Meas of Corr | Infor Meas of Corr 2 |
| Authentic | 3.23E-03 | 1.62E+01 | 8.25E-01 | 4.71E+01 | 2.76E-01 | 2.60E+02 | 1.72E+02 | 5.74E+00 | 8.76E+00 | 7.71E-04 | 3.14E+00 | -1.77E-01 | 8.97E-01 |
| FILM_1 | 5.63E-03 | 7.47E+00 | 8.90E-01 | 3.48E+01 | 3.77E-01 | 2.59E+02 | 1.32E+02 | 5.53E+00 | 7.99E+00 | 1.20E-03 | 2.62E+00 | -2.55E-01 | 9.45E-01 |
| FILM_3 | 5.48E-03 | 8.03E+00 | 8.83E-01 | 3.49E+01 | 3.70E-01 | 2.61E+02 | 1.32E+02 | 5.54E+00 | 8.05E+00 | 1.14E-03 | 2.67E+00 | -2.45E-01 | 9.41E-01 |
| FILM_7 | 5.02E-03 | 8.80E+00 | 8.83E-01 | 3.77E+01 | 3.61E-01 | 2.62E+02 | 1.42E+02 | 5.61E+00 | 8.18E+00 | 1.10E-03 | 2.73E+00 | -2.42E-01 | 9.42E-01 |
| Linear_1 | 5.00E-03 | 9.25E+00 | 8.61E-01 | 3.40E+01 | 3.38E-01 | 2.60E+02 | 1.27E+02 | 5.51E+00 | 8.13E+00 | 1.07E-03 | 2.77E+00 | -2.17E-01 | 9.23E-01 |
| Linear_3 | 6.40E-03 | 9.63E+00 | 8.52E-01 | 3.31E+01 | 3.39E-01 | 2.62E+02 | 1.23E+02 | 5.47E+00 | 7.95E+00 | 1.04E-03 | 2.79E+00 | -2.14E-01 | 9.17E-01 |
| Linear_7 | 5.33E-03 | 1.03E+01 | 8.54E-01 | 3.52E+01 | 3.30E-01 | 2.62E+02 | 1.30E+02 | 5.54E+00 | 8.14E+00 | 1.01E-03 | 2.83E+00 | -2.12E-01 | 9.20E-01 |

## Notes

**Conflict of interest statement** The authors declare no conflicts of interest.

### Competing Interest Statement

The authors have declared no competing interest.

### Summary of Updates

THis version includes new supplemental videos of interpolated datasets; and grant acknowledgment.

